# *Shigella flexneri* disruption of host cell-cell tension promotes intercellular spread

**DOI:** 10.1101/790147

**Authors:** Jeffrey K. Duncan, Alexandra L. Wiscovitch, Marcia B. Goldberg, Brian C. Russo

**Affiliations:** Division of Infectious Diseases, Department of Medicine, Massachusetts General Hospital, Boston, MA 02114, USA; Research Scholar Initiative, The Graduate School of Arts and Sciences, Harvard University, Cambridge, MA 02138, USA; Department of Microbiology, Blavatnik Institute, Harvard Medical School, Boston, MA 02115, USA

**Keywords:** *Shigella flexneri*, intercellular spread, intracellular pathogens, type 3 secretion system, plasma membrane

## Abstract

During infection, a subset of bacterial pathogens invades into the eukaryotic cytosol and spreads between cells of an epithelial layer. This intercellular spread is essential for disease and requires actin-based motility leading to the formation of plasma membrane protrusions. Protrusions are engulfed by the adjacent cell in an active process requiring both bacterial and eukaryotic proteins. Here, we demonstrate that the *Shigella* spp. type 3 secretion system protein IpaC promotes bacterial spread by reducing intercellular tension. *S. flexneri* producing a point mutant of IpaC that cannot interact with the cell-cell adhesion protein β-catenin were unable to reduce intercellular tension, form protrusions, or spread, demonstrating that interaction of IpaC with β-catenin is required for these processes. Spread was restored by chemical reduction of intercellular tension or genetic depletion of β-catenin. This work defines a molecular mechanism by which *Shigella* overcomes host cell-cell tension to mediate spread.

## Introduction

Many cytosol-dwelling bacterial pathogens have evolved unique mechanisms of spreading directly from the cytosol of an infected (donor) cell into an uninfected, adjacent (recipient) cell. Spreading enables the pathogens to avoid immune clearance, access new nutrients, and maintain an intracellular lifecycle (Sansonetti et al., 1991; Weddle and Agaisse, 2018a). *Shigella flexneri*, a model cytosol-dwelling Gram-negative bacterial pathogen, infects the colonic epithelium, replicating within the cytosol of colonocytes and subsequently spreading through the epithelium (Labrec et al., 1964; Sansonetti et al., 1986). Intercellular spread is required for *S. flexneri* to cause disease and efficiently colonize the colon (Sansonetti et al., 1991; Yum et al., 2019). *S. flexneri* polymerizes actin and moves in a directed manner to the cell periphery (Bernardini et al., 1989; Egile et al., 1999; Goldberg and Theriot, 1995), where it remodels the plasma membrane into pathogen-containing protrusions (Kadurugamuwa et al., 1991; Robbins et al., 1999). These bacteria-containing protrusions are engulfed into a vacuole by recipient cells in a clathrin-dependent process (Fukumatsu et al., 2012). Bacteria escape this vacuole into the cytosol of the donor cell (Allaoui et al., 1992; Campbell-Valois et al., 2015; Uchiya et al., 1995; Weddle and Agaisse, 2018b), enabling repeated cycles of intercellular spread through the epithelium. Whereas the forces derived from actin-based motility are necessary for protrusion formation (Monack and Theriot, 2001), other pathogen and host factors, including the type 3 secretion translocon pore proteins IpaB and IpaC and the type 3 effectors OspE1/2, IcsB, and VirA, are required for efficient intercellular spread (Allaoui et al., 1995; Campbell-Valois et al., 2015; Campbell-Valois et al., 2014; Heindl et al., 2010; Kuehl et al., 2014; Ogawa et al., 2003; Page et al., 1999; Schuch et al., 1999; Yi et al., 2014; Yoshida et al., 2006). Understanding the molecular mechanisms by which these proteins contribute to spread will define the parameters necessary for bacterial spread.

Here, we show that the intercellular spread of *S. flexneri* is dependent upon IpaC. We show that IpaC binds to the cell-cell adhesion protein β-catenin and that the IpaC-β-catenin interaction enables intercellular spread by overcoming cell-cell tension; overcoming cell-cell tension is required for the bacteria to generate protrusions of the plasma membrane. Moreover, substitution of an arginine residue in the C-terminal tail of IpaC abrogates the interaction of IpaC with β-catenin, consequently diminishing protrusion formation and spread. Our results define a mechanism by which *S. flexneri* subverts cell processes at the plasma membrane, enabling its spread between cells.

## Results

### The IpaC C-terminal Tail is Required for Efficient Intercellular Spread of *S. flexneri*

During invasion by *S. flexneri*, the type 3 secreted protein IpaC interacts with intermediate filaments; this interaction is necessary for efficient bacterial docking onto host cells and for efficient translocation of effectors into the host cell cytosol, where the effectors mediate bacterial invasion (Russo et al., 2016). In addition to being required for invasion, type 3 secretion is required for *S. flexneri* intercellular spread (Campbell-Valois et al., 2014; Page et al., 1999), and we hypothesized that the interaction between IpaC and intermediate filaments is required at this step. We tested the efficiency of spread for *S. flexneri* Δ*ipaC* producing WT IpaC or an IpaC derivative that contains a point mutation near the C-terminus (IpaC R362W) and is unable to interact with intermediate filaments (Harrington et al., 2006; Russo et al., 2016; Terry et al., 2008). IpaC R362W is efficiently secreted and forms normal-sized pores in the plasma membrane during invasion (Russo et al., 2019; Russo et al., 2016). Bacterial plaques formed in monolayers of mouse embryonic fibroblasts (MEFs) were significantly smaller for *S. flexneri* Δ*ipaC* producing IpaC R362W than for *S. flexneri* Δ*ipaC* producing WT IpaC (Figure 1A-B). In contrast, the presence of intermediate filaments did not affect plaque size (Figure 1A-B). Similarly, plaques formed by wild-type *S. flexneri* in monolayers of MEFs lacking vimentin (Vim^−/-^) were similar to those formed by MEFs producing vimentin (Vim^+/+^) (Figure 1C); vimentin is the only intermediate filament in these cells (Colucci-Guyon et al., 1994; Holwell et al., 1997), indicating that other intermediate filaments are not compensating for the loss of vimentin. Together these data show that the C-terminal region of IpaC at Arginine 362 is required for spread independent of the intermediate filament vimentin.

**Figure 1.**
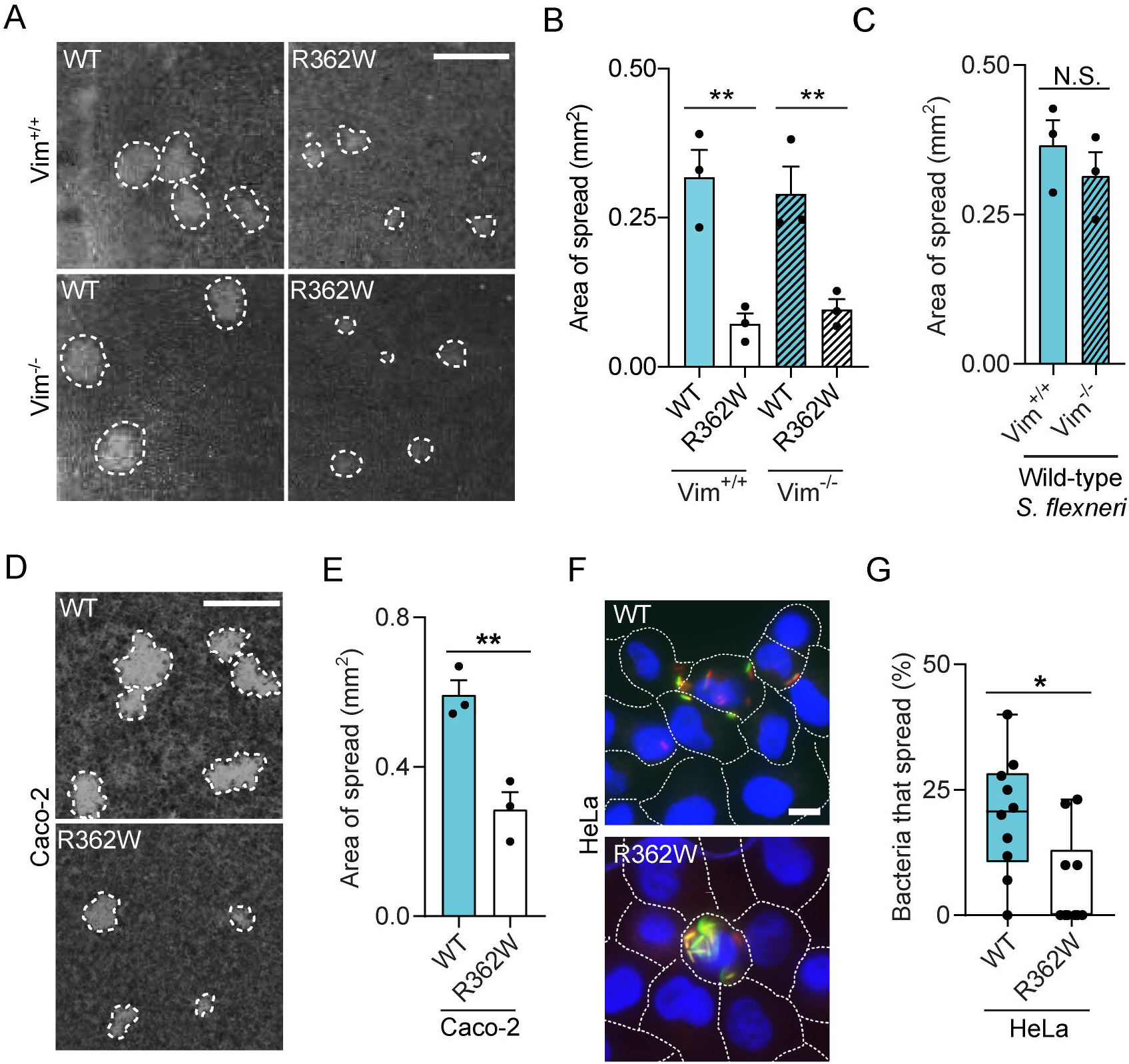
IpaC C-terminal tail is required for efficient intercellular spread of *S. flexneri*. (A-B) Plaques formed in cell monolayers by *S. flexneri* Δ*ipaC* producing wild-type IpaC or IpaC R362W. Representative Images. (B) Plaque size from experiments represented in A. (C) Plaques formed by wild-type *S. flexneri* in Vim^+/+^ and Vim^−/-^ MEFs. Representative images. (D) Plaques formed by *S. flexneri* strains in Caco-2 monolayers. Representative images. (E) Plaque size from experiments represented in panel D, (F) Infectious foci formed in HeLa cells by *S. flexneri* strains. Red, *S. flexneri*; green, bacteria with active T3SS; blue, DAPI. Representative images. (G) Percentage of bacteria that escape into adjacent cells from experiments represented in panel F. For box plots, Dots represent independent experiments (B, C, and E); data are mean ± SEM. Dots represent individual infectious foci of one experiment from two independent experimental replicates (G); boxes outline the 25th and 75th percentiles, midlines denote medians, and whiskers show minimum and maximum values. Scale bars, 500 µm (A, D) or 10 µm (F). N.S., not significant; *, p<0.05; **, p<0.01.

*S. flexneri* infects epithelial cells of the intestine, in which the predominant intermediate filaments are keratins rather than vimentin. Therefore, the ability of *S. flexneri* to spread was tested in Caco-2 cells, which express keratin intermediate filaments. Similar to the results with MEFs, plaques formed in monolayers of Caco-2 cells were significantly smaller for *S. flexneri* Δ*ipaC* producing IpaC R362W than for *S. flexneri* Δ*ipaC* producing WT IpaC (Figure 1D-E). The defect was again independent of intermediate filaments, as wild type *S. flexneri* formed plaques of similar size in Caco-2 cells whether or not the intermediate filaments keratin 8 or keratin 18 were present (Figure S1). Altogether, these data show the spread of *S. flexneri* is independent of intermediate filaments and dependent upon the presence of the IpaC C-terminal residue R362.

To distinguish whether IpaC aids in intercellular spread by contributing to a bacterial process in the donor cell or the recipient cell, we directly assessed the efficiency of *S. flexneri* spread from the initial donor cell to adjacent recipient cells. To do so, a population of cells was sparsely seeded and infected. Uninfected cells were overlaid onto the infected cells, and the ability of *S. flexneri* to spread into the second population of cells was assessed at 4 hours of infection. Compared to bacteria producing WT IpaC, bacteria producing IpaC R362W were impaired in movement out of the donor cells (Figure 1F-G), demonstrating that IpaC, and specifically residue R362 adjacent to the C-terminus, is required during exit from donor cells.

### IpaC is Necessary for Efficient Formation of Plasma Membrane Protrusions

To ascertain the step of cellular exit at which IpaC R362W is defective, we assessed the efficiency of *S. flexneri* Δ*ipaC* producing IpaC variants to form plasma membrane protrusions. In HeLa cells, the percentage of intracellular bacteria located in membrane protrusions was markedly decreased for bacteria producing IpaC R362W compared to bacteria producing wild-type IpaC (Figure 2A–B), which demonstrates that IpaC is required for the formation and/or maintenance of bacterial protrusions of the membrane and that IpaC R362W is defective in this process.

**Figure 2.**
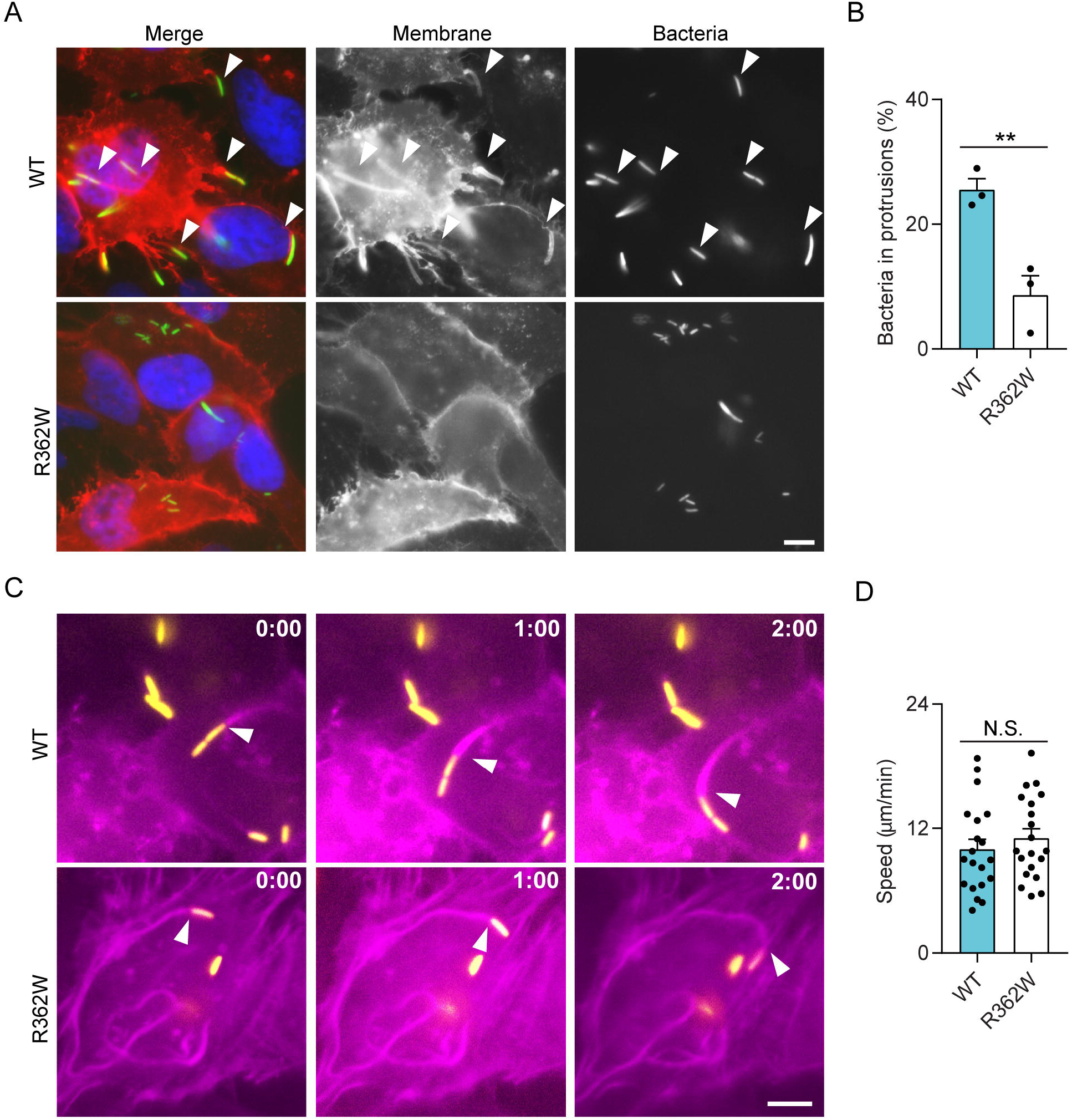
IpaC is required for protrusion formation by *S. flexneri*. Infection of HeLa cells with *S. flexneri* Δ*ipaC* producing wild-type IpaC or IpaC R362W. (A) Formation of plasma membrane protrusions. Green, *S. flexneri*; red, t-farnesyl-red fluorescent protein (RFP), which labels plasma membranes; blue, DAPI. Arrowheads, bacteria in protrusions. Representative images. (B) Percentage of intracellular bacteria located within protrusions from experiments represented in panel A, mean ± SEM. (C) Live cell imaging snapshots of actin-based motility of designated *S. flexneri* strains. Yellow, *S. flexneri*; purple, actin. Arrowheads, motile bacteria with unipolar polymerized actin. Representative images. (D) Speeds of bacteria with polymerized actin at one pole. Data is from two (C and D) or three (A and B) independent experiments. Dots represent independent experiments (B) or individual bacteria (D). Scale bars, 10 µM. N.S., not significant; **, p<0.01.

Given that forces derived from actin-based motility are required for and under heterologous experimental conditions can be sufficient for the formation of membrane protrusions (Makino et al., 1986; Monack and Theriot, 2001), we hypothesized that bacteria producing IpaC R362W might be defective in actin-based motility. To evaluate the efficiency of actin polymerization, we measured the speed of intracellular bacteria with actin tails using live microscopy in HeLa cells stably producing GFP-tagged actin (LifeAct). The speeds were similar for bacteria producing IpaC R362W and bacteria producing wild-type IpaC (Figure 2C–D), which indicates that IpaC plays a role in protrusion formation in a manner independent of actin-based motility.

In support of the defect of IpaC R362W-producing bacteria occurring at the stage of protrusion formation *per se*, we found no defect in other post-entry processes (Figure S2). Bacteria producing IpaC R362W efficiently escaped from the vacuole into the cytosol (Figure S2B). They regulated effector secretion through the T3SS similar to WT IpaC both in the timing of effector secretion (Figure S2C-D) and in the magnitude of secretion activation (Figure S2C and S2E). Unlike an *icsB* mutant, they avoided recruitment of autophagy components similar to bacteria producing WT IpaC (Figure S2F). Moreover, the percentage of bacteria that assembled an actin tail was similar to WT IpaC (Figure S2G-H), indicating that the reduction in protrusion formation (Figure 2A-B and Figure S2I) was not due to a defect in initiation of actin tail formation by IpaC R362W-producing bacteria. Altogether these data indicate that IpaC is required for formation of plasma membrane protrusions *per se* and that *S. flexneri* Δ*ipaC* producing IpaC R362W is impaired in spread due to a defect in its ability to form protrusions.

### Cell-cell Tension is Reduced by *S. flexneri* in an IpaC-Dependent Manner

*Listeria monocytogenes* and *Rickettsia parkeri* reduce cell-cell tension during spread. The secreted proteins Internalin C (*L. monocytogene*) and Sca4 (*R. parkeri*) (Lamason et al., 2016; Rajabian et al., 2009) decrease cell-cell tension, enabling the formation or resolution of plasma membrane protrusions, respectively. We investigated whether IpaC may similarly decrease intercellular tension. The impact of *S. flexneri* infection on the junctional linearity of the plasma membrane was measured (Otani et al., 2006). Whereas cells with normal membrane tension have nearly straight membranes between adjacent cell-cell vertices, cells that have decreased cell-cell membrane tension contain membranes that are more curved and thus have longer lengths of membrane between the vertices. Caco-2 cells infected with *S. flexneri* Δ*ipaC* producing wild-type IpaC displayed membranes that curved more than membranes of either cells infected with *S. flexneri* Δ*ipaC* producing IpaC R362W or uninfected cells (Figure 3A–B), indicating that, during infection, *S. flexneri* decreases cell-cell tension in an IpaC-dependent fashion.

**Figure 3.**
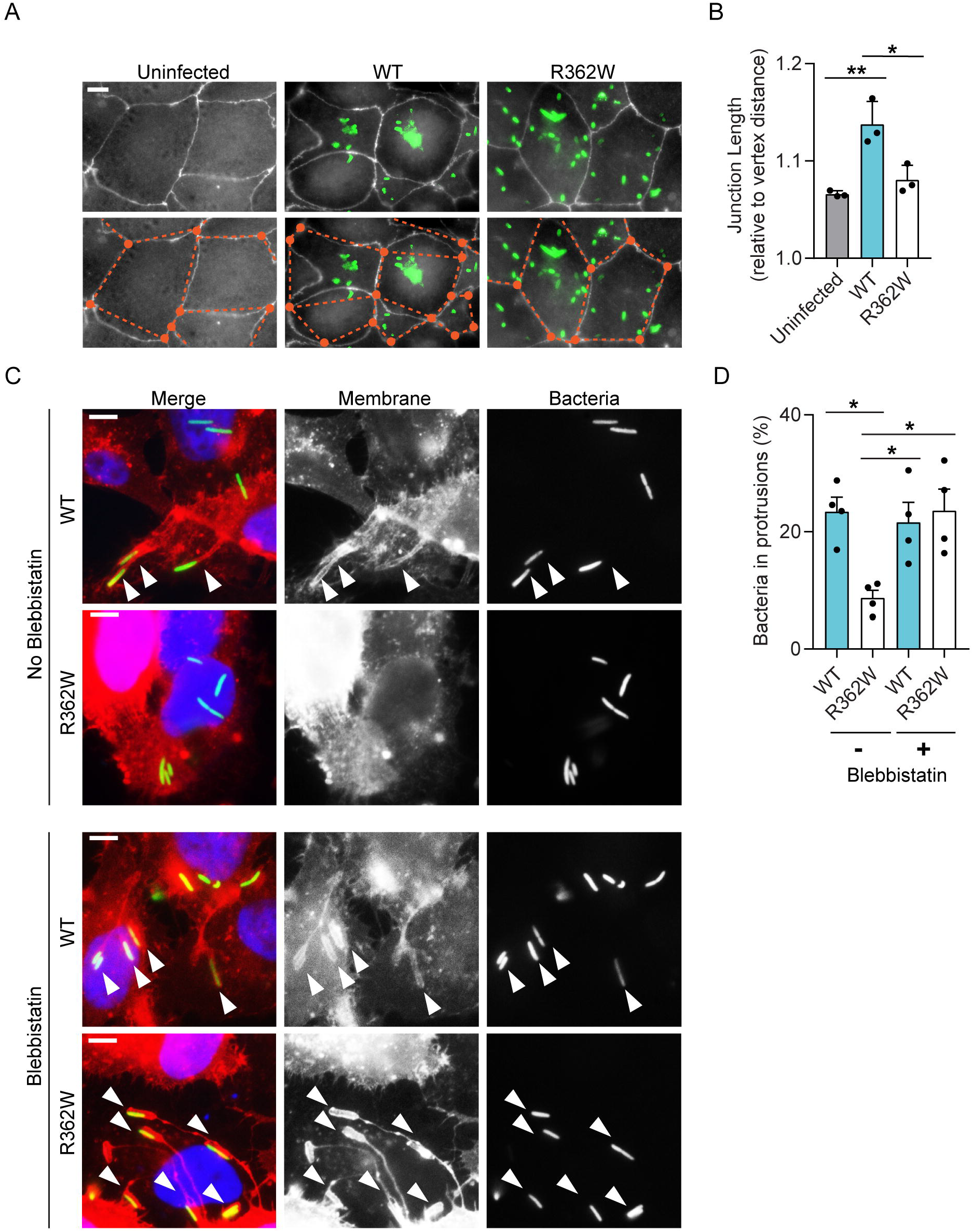
IpaC C-terminal tail arginine is required for *S. flexneri*-mediated reduction of host membrane tension. Infection of polarized Caco-2 cells with *S. flexneri* Δ*ipaC* producing wild-type IpaC or IpaC R362W. (A) Cell-cell junctions of Caco-2 cells delineated by ZO-1 staining. White, ZO-1; green, *S. flexneri*; orange dots, membrane junctions; orange dotted lines, linear distances between junctions. Representative images. (B) Membrane length from experiments represented in panel A, mean ± SEM. (C) Formation of plasma membrane protrusions after treatment with blebbistatin. Arrowheads, bacteria in protrusions. Red, t-farnesyl-RFP; green, *S. flexneri*; blue, DAPI. Representative Images. (D) Percentage of bacteria in protrusions from experiments represented in panel C, mean ± SEM. Dots represent independent experiments (B and D). Scale bars,10 µM. *, p<0.05; **, p<0.01.

Since the cortical actomyosin network maintains cell-cell tension at the adherens junctions, we tested whether IpaC reduces actomyosin-mediated tension in a manner that promotes protrusion formation. We assessed protrusion formation in the presence of blebbistatin, a chemical inhibitor of myosin II; inhibition of myosin II relieves cell-cell tension (Lamason et al., 2016; Rajabian et al., 2009). Blebbistatin treatment fully rescued protrusion formation of *S. flexneri* that produce IpaC R362W (Figure 3C–D), whereas it had no effect on the efficiency of protrusion formation by bacteria that produce wild-type IpaC, demonstrating that efficient formation of protrusions requires a reduction in actomyosin-mediated membrane tension.

### *S. flexneri* spread requires an interaction of IpaC with β-catenin

Catenin-cadherin networks maintain cell-cell tension and are integral to protecting against membrane stress (Ray et al., 2013). Since IpaC interacts with the cell-cell adhesion protein β-catenin (Shaikh et al., 2003), a component of these catenin-cadherin networks, we hypothesized that the role of IpaC in disruption of cell-cell tension may depend on its interaction with β-catenin and that IpaC R362 may be required for this interaction. In an orthogonal yeast-based protein-protein interaction assay that has been successfully used in the study of host-pathogen protein interactions (de Groot et al., 2011; Russo et al., 2016; Schmitz et al., 2009; Yi et al., 2014), we observed that IpaC R362W is defective for interaction with β-catenin. β-catenin (bait) was fused to the reovirus scaffold protein µNS, which forms inclusions bodies, and IpaC derivatives (prey) were tagged with mCherry. Protein-protein interaction results in the formation of puncta of fluorescent signal, whereas lack of interaction between bait and prey proteins results in diffuse mCherry signal throughout the cell (Figure 4A). Co-expression of IpaC-mCherry and β-catenin fused to µNS resulted in fluorescent foci in most yeast, whereas co-expression of IpaC R362W-mCherry and β-catenin fused to µNS resulted diffuse mCherry signal with few yeast displaying foci (Figure 4B–C). These data confirm that IpaC interacts with β-catenin and demonstrate that the IpaC R362W point mutation disrupts it.

**Figure 4.**
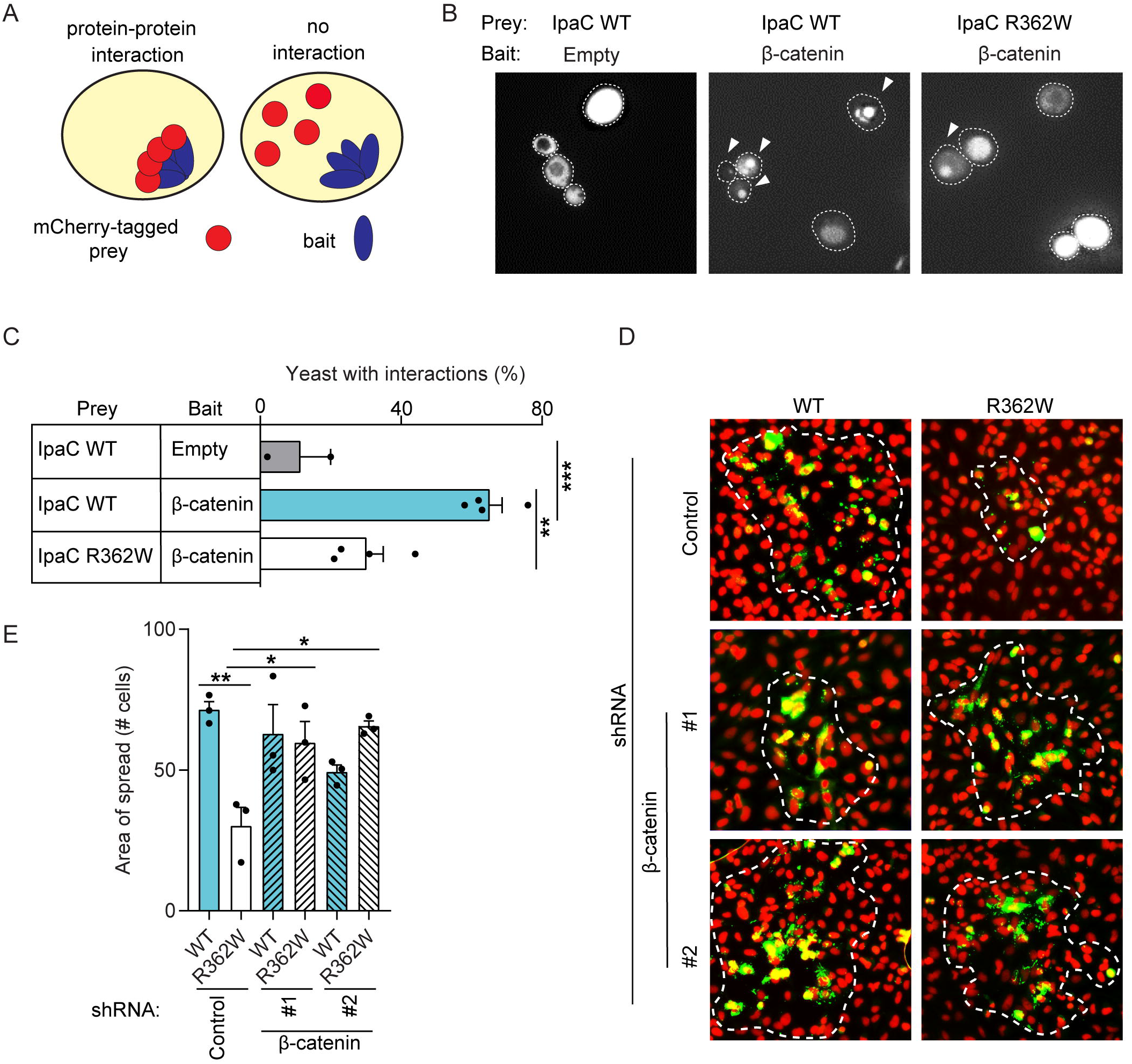
IpaC interactions with β-catenin are associated with *S. flexneri* cell to cell spread. (A) Schematic of yeast protein-protein interaction platform. Interaction of the mCherry-tagged prey protein (red) with the bait protein fused to the inclusion-body forming protein µNS (blue) results in puncta of red fluorescence. In contrast, lack of interaction between prey and bait proteins results in diffuse mCherry fluorescence throughout the cytosol of the yeast cells. (B) Protein interaction assay. Arrowheads, fluorescent puncta. Representative images. (C) Percentage of yeast displaying puncta indicating an interaction, mean ± SEM. (D) Bacterial plaques formed in monolayers stably expressing β-catenin-targeting (#1, #2) or control shRNA. Infection with *S. flexneri* Δ*ipaC* producing wild-type IpaC or IpaC R362W. Green, *S. flexneri*; blue, DAPI. Scale bar, 50 mm. Representative images. (E) Quantification of plaque size (area of spread) from experiments represented in panel D. Dots represent data from three or more independent experiments (C and E). *, p<0.05; **, p<0.01; *** p<0.001.

We tested the impact of β-catenin depletion on the efficiency of *S. flexneri* spread. Stable knock down of β-catenin in HeLa cells and Caco-2 cells (Figure S3A–D) rescued the spread of *S. flexneri* Δ*ipaC* producing IpaC R362W (Figures 4D-E and S3E). As expected, in cells expressing a scrambled shRNA, *S. flexneri* Δ*ipaC* producing IpaC R362W spread less efficiently than bacteria producing wild-type IpaC (Figure 4D–E). Knockdown of β-catenin on its own was insufficient to alter membrane linearity (Figure S3F), indicating that additional stresses associated with bacterial infection or other bacterial proteins are likely required to reduce membrane tension. These data demonstrate that β-catenin is a negative regulator of *S. flexneri* intercellular spread and that the interaction of IpaC with β-catenin disrupts β-catenin function in membrane tension.

## Discussion

For cytosol-dwelling bacterial pathogens, direct spread from an infected cell into adjacent uninfected cells is essential for dissemination during infection and disease pathogenesis. To traverse the plasma membranes, bacteria remodel the cellular cortical cytoskeleton. Here, we show that the *S. flexneri* protein IpaC decreases cell-cell tension and promotes the formation of plasma membrane protrusions, enabling intercellular spread, and that cell-cell tension was alleviated by the binding of IpaC to the cell-cell adhesion protein β-catenin.

β-catenin and cadherins are components of adherens junctions, which function to anchor one cell to another. β-catenin contains a stretch of 12 armadillo repeats. IpaC binds β-catenin within this armadillo repeat region, and its binding requires the ninth armadillo repeat (Figure S4) (Shaikh et al., 2003). E-cadherin binds via its cytoplasmic domain to β-catenin; in a co-crystal structure (PDB:1I7X), the interaction of E-cadherin with β-catenin extends across all 12 armadillo repeats (Figure S4) (Huber and Weis, 2001). Thus, the site of IpaC binding to β-catenin overlaps with the region of β-catenin to which E-cadherin binds (Figure S4), suggesting that IpaC may alter the interaction of E-cadherin and β-catenin. We speculate that IpaC disruption of cell-cell tension is mediated at least in part by modulation of the E-cadherin interaction with β-catenin.

Our data highlight that for efficient spread, *S. flexneri* must simultaneously maintain cell-cell contact but release membrane tension. Previous work shows cell-cell contact is necessary for intercellular spread; cell-cell adhesion proteins, including cadherin, tricellulin and occludin were required for efficient intercellular spread by *S. flexneri* (Fukumatsu et al., 2012; Sansonetti et al., 1994). In contrast, our findings show that the cell junction protein β-catenin restricts intercellular spread. Our findings that β-catenin restricts bacterial spread appear to oppose the role of E-cadherin in promoting spread. A model to explain these observations is that the interaction of IpaC with β-catenin may generate slack in cell-cell junctions while maintaining sufficient homotypic interactions of E-cadherin molecules to maintain cell-cell contact. Thus, by targeting catenin-cadherin interactions via β-catenin, IpaC may selectively alter membrane tension while maintaining cell-cell contact.

Our findings demonstrate that at different stages of *S. flexneri* infection, the arginine immediately adjacent to the C-terminus of IpaC (R362) is required for interactions with two distinct cellular proteins. In addition to being required for the formation of protrusions during intercellular spread, IpaC R362 is required for interactions with intermediate filaments to support docking during bacterial entry (Russo et al., 2016). Whether the binding preference of IpaC is determined by the distinct sub-cellular niche of the bacterium at these two stages of infection or by other factors is at present unclear. Of note, other *S. flexneri* type 3 secreted proteins contain sequences that bind more than one host protein, for example, during entry, vinculin binding site 3 of IpaA separately binds talin and vinculin (Valencia-Gallardo et al., 2019).

Like *S. flexneri*, *R. parkeri* and *L. monocytogenes* diminish cell-cell tension but do so by targeting different host proteins. *R. parkeri* prevents vinculin-mediated cell tension via the type 4 secretion system effector protein Sca4, whereas the secreted *L. monocytogenes* protein Internalin C decreases cell-cell tension by binding the focal adhesion protein Tuba (Lamason et al., 2016; Rajabian et al., 2009). These findings provide evidence of convergent evolution by a cadre of pathogens upon mechanisms to decrease cell-cell tension and underscore that cell-cell tension is a critical barrier that must be subverted for successful bacterial infection.

## Acknowledgements

We thank Ramnik Xavier, Victor Faundez, and Claude Parsot for reagents. We thank Cammie Lesser, Amy Barczak, Rebecca Lamason, and Allen Sanderlin for reagents and helpful discussions. We thank the members of the Goldberg, Lesser, and Barczak laboratories for helpful discussions. We thank Natasha Bitar for technical support and Brianna Lowey for helpful discussion and critical reading of the manuscript.

This work was funded by NIH grant AI081724 to M.B.G. and by NIH grants AI007061 and AI114162, the Massachusetts General Hospital Executive Committee on Research Tosteson Award, and the Charles A. King Trust Postdoctoral Research Fellowship Program, Bank of America, N.A., Co-Trustees, to B.C.R.

## Author Contributions

This project was conceived of and experiments were designed by J.K.D., M.B.G., and B.C.R. All experiments were performed by J.K.D., A.L.W., and B.C.R. The manuscript was written by J.K.D., A.L.W., M.B.G., and B.C.R.

## Supplemental Figure Legends

**Figure S1. Intermediate filaments keratin 8 and 18 are dispensable for intercellular spread of *S. flexneri.* (Related to Figure 1)**.

Plaque formation by wild-type *S. flexneri* in monolayers of Caco-2 cells stably expressing keratin 8 shRNA, keratin 18 shRNA, or non-targeting shRNA. (A) Representative images of plaques. (B) Plaque size (area of spread) from experiments represented in panel A, mean ± SEM. a.u., arbitrary units. Dots represent independent experiments.

**Figure S2. Vacuolar escape, regulation of the T3SS, autophagic escape, and actin tail formation of *S. flexneri* producing IpaC R362W are similar to that of *S. flexneri* producing wild-type IpaC. (Related to Figure 2)**.

Infections of cells with *S. flexneri* Δ*ipaC* producing wild-type IpaC or IpaC R362W. (A) Schematic of *S. flexneri* intracellular lifestyle. (I) After invasion, the *S. flexneri* T3SS ruptures the uptake vacuole, releasing bacteria into the cytosol. (II) In the cytosol, *S. flexneri* utilizes host actin machinery to propel itself to the periphery. (III) At the periphery, motile bacteria form plasma membrane protrusions. (IV) The membranous protrusion is engulfed into a double membrane vacuole in an adjacent cell. (V) The bacteria then rupture the secondary vacuole, thereby escaping into the cytosol, wherein the process is repeated. Type 3 secretion is active during invasion and during spread (stages I, III, IV). Green bacteria, active type 3 secretion. (B) Percent of bacteria that escape the vacuole of Vim^+/+^ MEFs, determined by chloroquine resistance. Intravacuolar bacteria are killed by the high vacuolar concentration of chloroquine, whereas bacteria that escape the vacuole into the cytosol survive. Mean ± SEM. (C) Vim^+/+^ MEFs infected for 60 or 150 minutes with bacteria containing TSAR, a fluorescent reporter of T3SS activity. Red, *S. flexneri*; green, *S. flexneri* with active T3SS; blue, DNA. Representative images. (D) Percent of *S. flexneri* with active T3SS from experiments represented in panel C and including additional time points at 90 and 120 minutes, mean ± SEM. (E) Intensity of the GFP signal from *S. flexneri* with active T3SS from panel C, mean ± SEM. (F) Percent of designated *S. flexneri* strains associated with GFP-LC3 during infection of GFP-LC3 HeLa cells. Positive control, *S. flexneri* ∆*icsB*. Mean ± SEM. (G) Actin tail and protrusion formation in Vim^+/+^ MEFs infected with designated *S. flexneri* strains for 180 minutes. Red, phalloidin: green, *S. flexneri*; blue, DAPI. Representative images. (H) Percentage of bacteria with unipolar actin (tails) from experiments represented in panel G, mean ± SEM. (I) Percentage of bacteria in protrusions from panel G, mean ± SEM. Dots represent independent experiments. Scale bars, 10 µM (C), 5 µM (G). N.S., not-significant; *, p<0.05; **, p<0.01.

**Figure S3. β-catenin in intercellular spread of *S. flexneri.* (Related to Figure 4)**.

(A) β-catenin levels in cell lysates from HeLa cells stably transfected with control or β-catenin targeting shRNAs. β-actin, loading control. Representative western blots. (B) Densitometric analysis of β-catenin depletion in HeLa cells from experiments represented in panel A. (C) β-catenin levels in cell lysates from Caco-2 cells stably transfected with control or β-catenin targeting shRNAs. β-actin, loading control. Representative western blots. (D) Densitometric analysis of β-catenin depletion in Caco-2 cells from experiments represented in panel C. (E) Plaque size (area of spread) of Caco-2 cells depleted of β-catenin and infected with *S. flexneri* Δ*ipaC* producing wild-type IpaC or IpaC R362W. (F) Quantification of membrane length for Caco-2 cells stably transfected with control or β-catenin targeting shRNAs. Mean ± SEM. Dots represent independent experiments. a.u., arbitrary units. * p<0.05; ** p<0.01; *** p<0.001.

**Figure S4. IpaC binding site on β-catenin overlaps with the E-cadherin binding pocket. (Related to Figures 1-4)**.

Region of β-catenin to which E-cadherin and IpaC bind. E-cadherin binds throughout the armadillo repeats (Huber and Weis, 2001). IpaC interaction with β-catenin occurs within the armadillo repeat region of β-catenin and requires the ninth armadillo repeat (Shaikh et al., 2003).

## STAR methods

### Contact for Reagent and Resource Sharing

Further information and requests for reagents may be directed to Marcia Goldberg or Brian Russo.

### Bacterial Culture

The wild-type *S. flexneri* strain used in this study is *S. flexneri* 2457T (Labrec et al., 1964), and all mutants were isogenic derivatives of it. *S. flexneri* strains were cultured in trypticase soy broth at 37°C. *ipaC* derivatives were cloned into the plasmid pBAD33, and their expression was driven from the pBAD promoter, induced with 1.2% arabinose.

### Cell Lines

HeLa, Caco-2, and MEFs were maintained at 37°C in 5% CO_2_. All cells were grown in Dulbecco’s modified Eagle’s medium (DMEM) supplemented with 10% fetal bovine serum (FBS). Caco-2 and HeLa cell lines stably expressing farnesylated TagRFP-T (gift of R. Lamason; HeLa only), shRNAs targeting β-catenin [Broad Institute; TRCN000314921 and TRCN000314991(Yang et al., 2011)], or a control (Addgene, Cat# 10879; (Moffat and Sabatini, 2006) were generated using retroviral transduction with selection with 10 μg/ml puromycin.

### Plaque Assays

For confluent monolayers, 8×10^5^ (Caco-2) or 6×10^5^ (MEF) cells per well were seeded in six-well plates. The next day, monolayers were infected with bacteria in mid-exponential phase at a multiplicity of infection (MOI) of 0.002 for MEFs and a MOI of 0.02 for Caco-2 cells in DMEM. Multiplicity of infection (MOI) was adjusted to compensate for the known decrease in efficiency of invasion by *S. flexneri* in the absence of interaction of IpaC with intermediate filaments. Bacteria were centrifuged onto cells at 800 x*g* for 10 minutes and incubated at 37°C in 5% CO_2_ for 50 minutes. Media was replaced with 0.5% agarose in DMEM containing 25 µg/mL gentamicin, 10% FBS, 1.2% arabinose (for *ipaC* expression from the arabinose promoter), and 0.45% glucose and incubated an additional two days. An additional overlay formulated as before but containing 0.7% agarose and 0.1% neutral red was added. Following incubation for at least 4h, the plates were imaged with an Epson Perfection 4990 photo scanner. ImageJ was used to quantify plaque area.

### Discrimination of primarily infected and secondarily infected cells

HeLa cells were seeded at 1×10^4^ cells per well on coverslips and, the next day, were infected at a MOI of 200, as above, with strains constitutively expressing a RFP reporter and a GFP whose expression is induced by T3SS secretion (pTSAR). At 30 min of infection, media was replaced with DMEM supplemented with 0.45% glucose, 1.2% arabinose, 25 µg/mL gentamicin, and 10% FBS. 6×10^4^ naïve HeLa cells were overlaid and centrifuged on onto the infected cells at 100 x*g* for 5 minutes. After an additional 3 hours, cells were fixed with 3.7% paraformaldehyde in phosphate-buffered saline (PBS) for 20 minutes and stained for 5 minutes with 1:10,000 Hoescht 33342 in PBS. Coverslips were mounted with Prolong Diamond Antifade Mountant (Thermo Fisher) and imaged the next day.

### Protrusion Assays

HeLa cells expressing t-farnesyl-RFP were seeded at 6×10^5^ cells per well on coverslips. The next day, cells were infected at an MOI of 100, as above, with designated strains expressing pROEX-Aqua [Addgene; plasmid #42889, (Erard et al., 2013)]. At 30 minutes of infection, media was replaced with DMEM supplemented with 0.45% glucose, 1.2% arabinose, 25 µg/mL gentamicin, 1 mM IPTG, and 10% FBS. After an additional 3 hours, cells were fixed with 3.7% paraformaldehyde in PBS and stained with for 5 minutes with 1:10,000 Hoescht 33342 in PBS. Coverslips were mounted with Prolong Diamond Antifade Mountant and imaged the next day.

### Live-cell Imaging Analysis

HeLa cells expressing *LifeAct-GFP* were seeded at 2×10^5^ cells per well on 20 mm MatTek glass bottom dishes. The next day cells were infected at a MOI of 100, as above, with *S. flexneri* expressing a constitutive RFP and a GFP reporter of T3SS activity (pTSAR). At 30 minutes of infection, media was replaced with DMEM supplemented with 0.45% glucose, 1.2% arabinose, 25 µg/mL gentamicin, 50 mM HEPES, and 10% FBS. Samples were imaged with a Nikon Eclipse TE-300 at 37°C at 5-second intervals for 5 minutes. To calculate actin-tail mediated velocities, the ImageJ plugin particle tracker was used to track in-frame bacteria over the course of the video. Speed was determined for each bacterium by measuring the distance traveled over the duration of time the bacteria was in focus.

### Membrane Linearity Assay

Membrane linearity was performed as previously described (Lamason et al., 2016). Caco-2 cells were seeded onto fibronectin treated coverslips at a density of 8×10^5^ cells per well and grown for 7 days to polarize the cells. After 4 days, the media was changed daily. Cells were infected as above with *S. flexneri* strains expressing the uropathogenic *E. coli* Afa-1 pilus (Labigne-Roussel et al., 1984), which binds to decay accelerating factor (CD55) on the surface of human cells (Nowicki et al., 1993) and enhanced infection at a MOI of 10. At 30 minutes of infection, media was replaced with DMEM, supplemented with 0.45% glucose, 1.2% arabinose, 25 µg/mL gentamicin, and 10% FBS. After an additional 3 hours, cells were fixed with 3.7% paraformaldehyde in PBS, permeabilized with 0.5% Triton X-100 in PBS, and stained with mouse anti-ZO-1 antibody (Invitrogen, cat# 339100; 1:100 dilution) and Alexa Fluor 568 conjugated secondary, with anti-*Shigella* antibody (ViroStat cat# 0903; 1:100 dilution), and with Hoescht 33342. Coverslips were mounted with Prolong Diamond Antifade Mountant and imaged the next day. The junction length was determined by measuring with ImageJ the linear distance between cell vertices and the actual length of the plasma membrane between the same vertices. Junction length was expressed as a ratio of the actual length of the membrane divided by the linear inter-vertex distance.

### Yeast Protein-Protein Interaction Assay

Yeast protein interaction assays were performed as previously described (Schmitz et al., 2009). Yeast carrying plasmids encoding mCherry-tagged IpaC variants and µNS alone (negative control) or µNS-tagged β-catenin were cultured overnight in complete synthetic media lacking histidine and leucine supplemented with 2% raffinose. The next morning, strains were back diluted to OD_600_ 0.5 and grown for 2h at 30°C with 2% raffinose. Then, to induce protein synthesis, the media was changed to 2% galactose, and growth was allowed to proceed for 4h at 30°C. Yeast were wet mounted and imaged. To determine the percentage of yeast with puncta, the total number of yeast cells was determined by brightfield and puncta of red fluorescent protein were counted.

### Infectious Foci Assay

HeLa cells were seeded at a density of 6 × 10^5^ cells per well on coverslips in 6 well plates. The next day, cells were infected as above at a MOI of 0.05 with designated strains. At 50 minutes of infection, media was replaced with DMEM supplemented with 0.45% glucose, 1.2% arabinose, 25 µg/mL gentamicin, and 10% FBS. Cells were incubated at 37°C with 5% CO_2_ for 12 hours at which point they were rinsed once and fixed with 3.7% paraformaldehyde in PBS for 20 minutes. Cells were washed three times with PBS, incubated with 1 M glycine in PBS for 15 minutes, washed three additional times with PBS and then permeabilized with 0.5% Triton X-100 for 20 minutes. Cells were washed five times with PBS and incubated overnight at 4°C with Alexa Fluor 488 conjugated anti-*Shigella* antibody (ViroStat, 1:1000 dilution). Cells were washed three times with PBS, stained with Hoescht 33342, washed twice with PBS and mounted with Prolong Diamond Antifade Mountant. Foci were randomly imaged across the coverslip, and the number of infected cells within each focus was determined by counting nuclei that co-localized with bacteria.

### Vacuolar Escape

The resistance of intracellular bacteria to chloroquine, which accumulates within vacuoles and kills intra-vacuolar bacteria but remains at sub-bactericidal concentrations within the cytosol, was tested as previously described (Zychlinsky et al., 1994). Notably, IpaC is an important factor in vacuolar escape ((Du et al., 2016), Briefly, 1×10^4^ Vim^+/+^ MEFs were seeded into wells of a 96-well plate. The next day, cells were infected as above at a MOI of 100 with designated strains. At 50 minutes of infection, media was replaced with DMEM supplemented with 0.45% glucose, 1.2% arabinose, 25 µg/mL gentamicin, 10% FBS ± 200 µg/mL chloroquine. After 1 hour, cells were washed three times and lysed with 1% Triton X-100 in PBS. Bacteria were serially diluted 1:5 and the number of intracellular bacteria were enumerated by plating dilutions. Percent cytosolic bacteria were the ratio of colony forming units (CFU) in the presence of gentamicin and chloroquine to CFU in the presence of gentamicin without chloroquine multiplied by 100.

### LC3 Co-localization

To quantify intracellular bacteria associated with LC3, HeLa cells stably expressing a GFP-LC3 construct were seeded at a density of 4 × 10^5^ cells per well in a 6-well plate on coverslips. The next day, cells were infected as above at a MOI of 100 with indicated strains. At 50 minutes of infection, media was replaced with DMEM supplemented with 0.45% glucose, 1.2% arabinose, 25 µg/mL gentamicin, and 10% FBS. After an additional 3 hours, cells were washed once with PBS, fixed with 3.7% paraformaldehyde and stained with Hoechst 33342. Coverslips were mounted with Prolong Diamond Antifade Mountant, and cells were imaged randomly across the coverslip. LC3-positive bacteria were counted as those that co-localized with strong GFP signal.

### Type 3 Secretion System Activity

To measure type 3 secretion system activity, Vim^+/+^ MEFs were seeded at a density of 4 × 10^5^ cells per well in 6-well plates on coverslips. The next day, cells were infected, as above, with indicated strains carrying the plasmid pTSAR [GFP expression is driven by the mxiE box, which requires type 3-mediated secretion of the effector OspD1, and mCherry is driven by the constitutive rpsM promoter (Campbell-Valois et al., 2014; Parsot et al., 2005). At 20 minutes of infection, media was replaced with DMEM supplemented with 0.45% glucose, 1.2% arabinose, 25 µg/mL gentamicin, and 10% FBS. At indicated time points, cells were washed once with PBS, fixed with 3.7% paraformaldehyde, and stained with Hoechst 33342. Cells were washed two additional times and mounted with Prolong Diamond Antifade Mountant. Cells were randomly imaged across the coverslip, and the percent of bacteria with active type 3 secretion systems was determined by enumerating the ratio of GFP-positive bacteria (T3SS active) to RFP-positive bacteria (total). To determine the intensity of GFP signal, bacteria were identified by an RFP signal, and the intensity of GFP signal was measured with imageJ (Schneider et al., 2012).

### Actin Tail Formation

To quantify actin-tail formation, Vim^+/+^ MEFs were seeded at a density of 4 × 10^5^ cells per well in 6-well plates on coverslips. The next day, cells were infected as above with indicated strains. At 50 minutes of infection, media was replaced with DMEM supplemented with 0.45% glucose, 1.2% arabinose, 25 µg/mL gentamicin, and 10% FBS. After an additional 3 hours, cells were washed once with PBS, fixed with 3.7% paraformaldehyde, and stained with anti-*Shigella* antibody conjugated to Alexa Fluor 488 (ViroStat, 1:1000 dilution). The next day, cells were washed three times with PBS and stained with phalloidin conjugated to Alexa Fluor 568 (Invitrogen, cat# A12380). Cells were washed three times with PBS and stained with Hoescht 33342. Cells were washed twice more with PBS and mounted with Prolong Diamond Antifade Mountant. The percentage of bacteria with actin-tails was determined by counting the number of total bacteria (GFP-positive) to the number of bacteria with actin-tails.

### Quantification and Statistical Analysis

Statistical differences between means was determined with GraphPad Prism. Statistical differences between two means was tested by unpaired Student’s t-test. Differences between the means of three or more groups was tested by either two-way ANOVA with Sidak *post hoc* test (Figures S2E-F) or one-way ANOVA with Tukey *post hoc* test. Microscopic images were pseudocolored and assembled using Adobe Photoshop or ImageJ.

