## Supplemental Figures for "*Shigella flexneri* disruption of host cell-cell tension promotes intercellular spread"

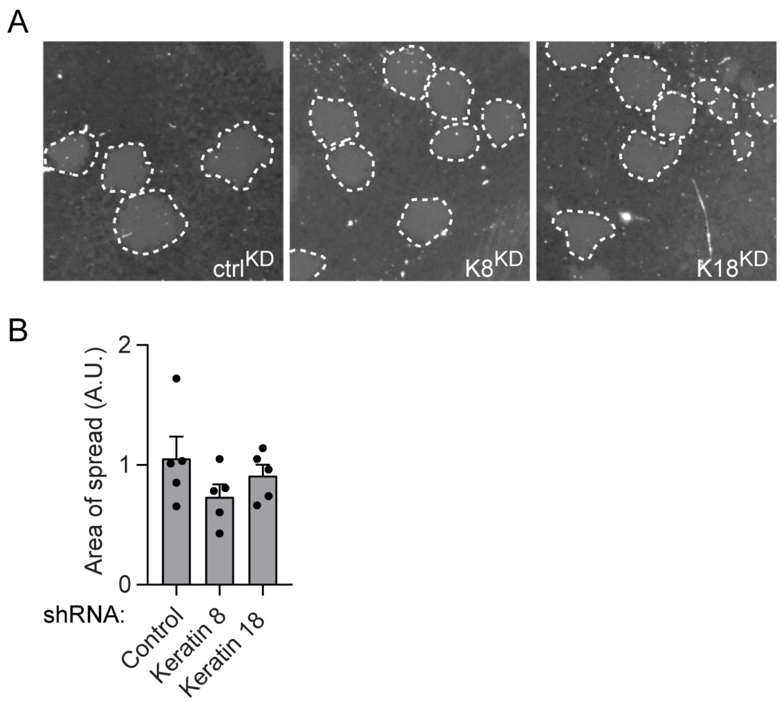


**Figure S1. Intermediate filaments keratin 8 and 18 are dispensable for intercellular spread of *S. flexneri.* (Related to Figure 1).**

Plaque formation by wild-type *S. flexneri* in monolayers of Caco-2 cells stably expressing keratin 8 shRNA, keratin 18 shRNA, or non-targeting shRNA. (A) Representative images of plaques. (B) Plaque size (area of spread) from experiments represented in panel A, mean ± SEM. a.u., arbitrary units. Dots represent independent experiments.

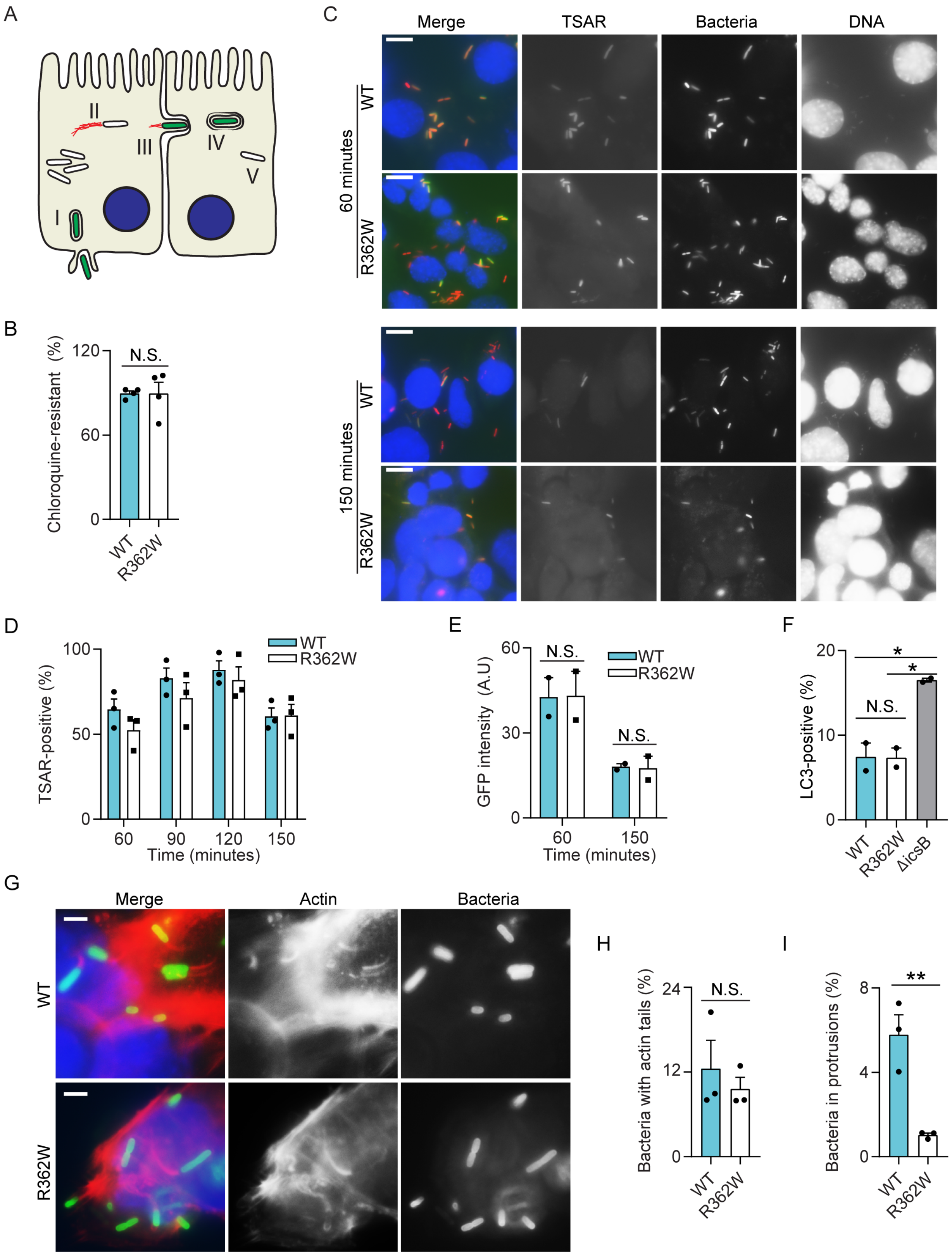
A
I
II
III
IV
V
B
Chloroquine-resistant (%)
N.S.
120
60
0
WT
R362W
C
Merge
TSAR
Bacteria
DNA
60 minutes
150 minutes
WT
R362W
D
TSAR-positive (%)
100
50
0
WT
R362W
60
90
120
150
Time (minutes)
E
GFP intensity (A.U)
N.S.
60
30
0
150
F
LC3-positive (%)
*
20
10
ΔicsB
G
Actin
H
Bacteria with actin tails (%)
24
12
I
Bacteria in protrusions (%)
**
8
4

**Figure S2. Vacuolar escape, regulation of the T3SS, autophagic escape, and actin tail formation of *S. flexneri* producing IpaC R362W are similar to that of *S. flexneri* producing wild-type IpaC. (Related to Figure 2).**

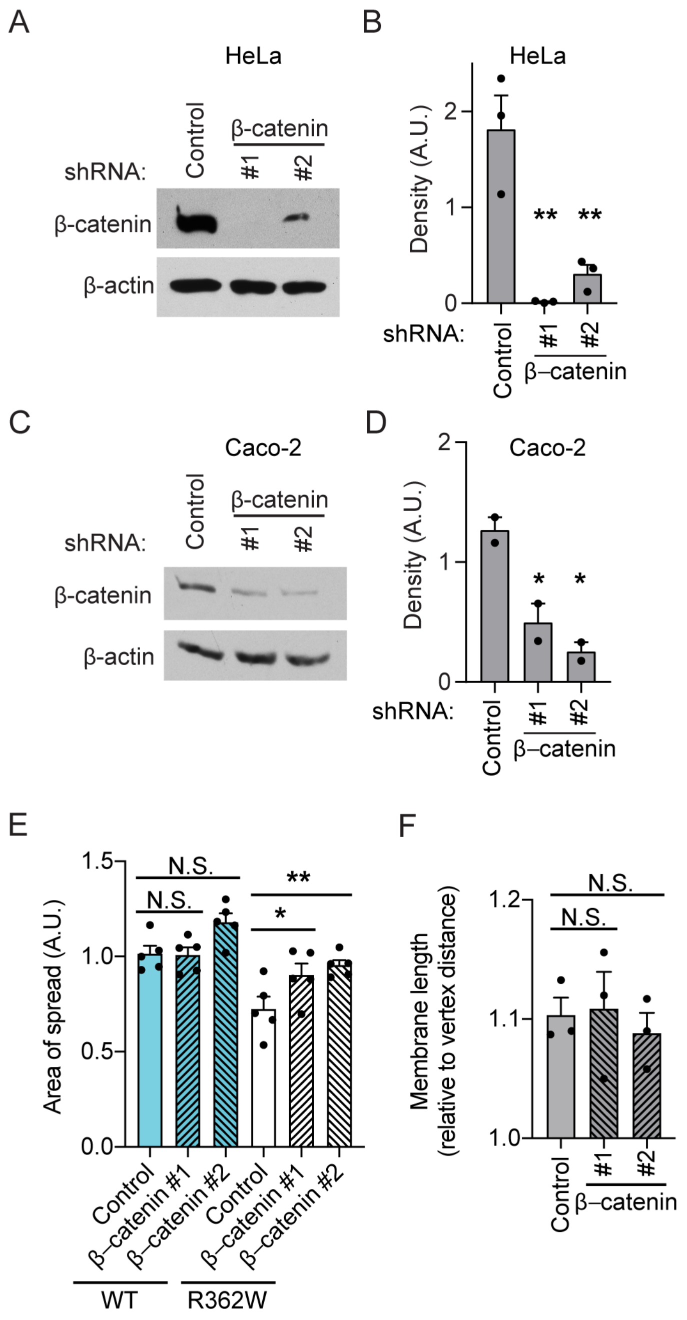


**Figure S3. β-catenin in intercellular spread of *S. flexneri.* (Related to Figure 4).**

(A) β-catenin levels in cell lysates from HeLa cells stably transfected with control or β-catenin targeting shRNAs. β-actin, loading control. Representative western blots. (B) Densitometric analysis of β-catenin depletion in HeLa cells from experiments represented in panel A. (C) β-catenin levels in cell lysates from Caco-2 cells stably transfected with control or β-catenin targeting shRNAs. β-actin, loading control. Representative western blots. (D) Densitometric analysis of β-catenin depletion in Caco-2 cells from experiments represented in panel C. (E) Plaque size (area of spread) of Caco-2 cells depleted of β-catenin and infected with *S. flexneri* ∆*ipaC* producing wild-type IpaC or IpaC R362W. (F) Quantification of membrane length for Caco-2 cells stably transfected with control or β-catenin targeting shRNAs. Mean ± SEM. Dots represent independent experiments. a.u., arbitrary units. * $p<0.05$; ** $p<0.01$; *** $p<0.001$.

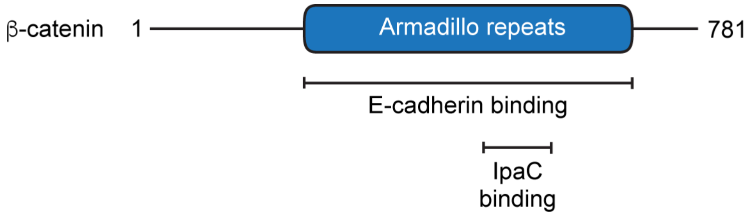


**Figure S4. IpaC binding site on β-catenin overlaps with the E-cadherin binding pocket. (Related to Figures 1-4).**

Region of β-catenin to which E-cadherin and IpaC bind. E-cadherin binds throughout the armadillo repeats (Huber and Weis, 2001). IpaC interaction with β-catenin occurs within the armadillo repeat region of β-catenin and requires the ninth armadillo repeat (Shaikh et al., 2003).
